# Engineering microbial consortia: uptake and leakage rate differentially shape community arrangement and composition

**DOI:** 10.1101/2024.07.19.604250

**Authors:** Estelle Pignon, Gábor Holló, Théodora Steiner, Simon van Vliet, Yolanda Schaerli

## Abstract

Bacteria often grow as communities in intricate spatial arrangements on surfaces and interact with each other through the local exchange of diffusible molecules. Yet, our understanding of how these interactions shape the properties of the communities remains limited. Here, we study synthetic communities of *Escherichia coli* amino acid auxotrophs interacting through the obligate exchange of amino acids. We genetically engineer these strains to alter their amino acid leakage and uptake abilities. We then characterise the spatial arrangement and composition of the communities when grown on a surface. By integrating experimental data with mathematical modeling, we demonstrate that amino acid uptake and leakage rates are crucial determinants of community structure. Our results show that while the spatial arrangement of the community is primarily governed by uptake rates, the community composition is predominantly influenced by leakage rates. These findings enhance our understanding of microbial community dynamics and provide a framework for predicting and engineering microbial consortia.

## Introduction

Microbial communities are ubiquitous on Earth and often grow on surfaces, for example in the form of biofilms [1]. Growth of surface-associated microbes is dependent on the physio-chemical properties of their local environment such as gradients of glucose, pH, viscosity or oxygen that influence the spatial arrangement of microbial communities [2–6]. In return, microbes actively change their environment as well, by consuming and releasing metabolites [7–9] or changing the local chemical properties [10]. Finally, microbes interact with each other, often through the exchange of diffusible molecules [11, 12]. On one hand, microbes leak a large diversity of metabolites [13], on the other hand, many microorganisms rely on the uptake of metabolites released by other community members. Indeed, many microbes are auxotrophic for certain compounds, meaning they are unable to synthesize a particular organic compound required for their growth [14]. This leads to diffusion-dominated interactions taking place in highly structured communities, making the microbial arrangement in space a crucial parameter for access to nutrients or the exchange of metabolites. Among the exchanged molecules, amino acids are highly prevalent [11, 14, 15] and amino acid cross-feeding can be found in a variety of environments, such as the mammalian gut [16, 17], the plant leaf [18] and even in oil reservoirs [19]. Amino acid auxotrophies have been shown to create strong interdependencies, which affect the dynamics of community compositions, carbon and energy fluxes, and host-microbiome interactions [20, 21]. They also promote high species diversity, drug tolerance, evenness and robustness of the consortium [22–24].

To study molecular interactions and the organisation of spatially structured microbial communities, range expansion assays are a powerful approach [25, 26]. They consist of inoculating either a single strain or a community, often fluorescently labelled, on an agar plate. The cells then expand radially, enabling the observation of spatial pattern formation. In addition, external parameters such as temperature, carbon sources, or inducer molecules can easily be modified. This setup enables one to study and understand in details the effect of a multitude of parameters on microbial communities such as mutualism and competition [27–29], trophic dependencies [6], short- or long-range weapons [30, 31], cellular growth rates [32, 33] or cell motility [34, 35].

Despite these recent advances, we still do not fully understand how molecular interactions between microbes determine the properties of communities, such as their composition and spatial arrangement. To address this question, Dal Co and colleagues studied a synthetic community of two auxotrophic *Escherichia coli* strains [36]. Each strain is unable to produce one essential amino acid (either tryptophan or proline), but they can grow together in a co-culture by exchanging these metabolites. The authors measured single-cell growth rates in a microfluidic device and concluded that the bacteria only interact with nearby cells. In a follow-up study, van Vliet and colleagues developed a mathematical framework [37] that derives community-level properties from the interactions at the local scale. In particular, the authors postulated that a few key biophysical parameters are sufficient to predict the arrangement and composition of the community. Those parameters are the cell density, as well as the leakage and uptake rates of the exchanged metabolites (i.e. amino acids in this case) (Figure 1). In a dense community, the spatial arrangement is predicted to be determined mostly by the uptake rate of the amino acids. Briefly, high uptake rates prevent the exchanged molecules from diffusing far, while low uptake rates create larger regions where the exchanged molecules are present. Hence, cells with a high uptake rate can only grow close to their partner, while a low uptake rate leads to growth further away from the partner and to bigger patches. In contrast, the composition of the community is predicted to be mainly determined by the leakage rates, with uptake rates mostly having a modulating effect [37].

**Figure 1:**
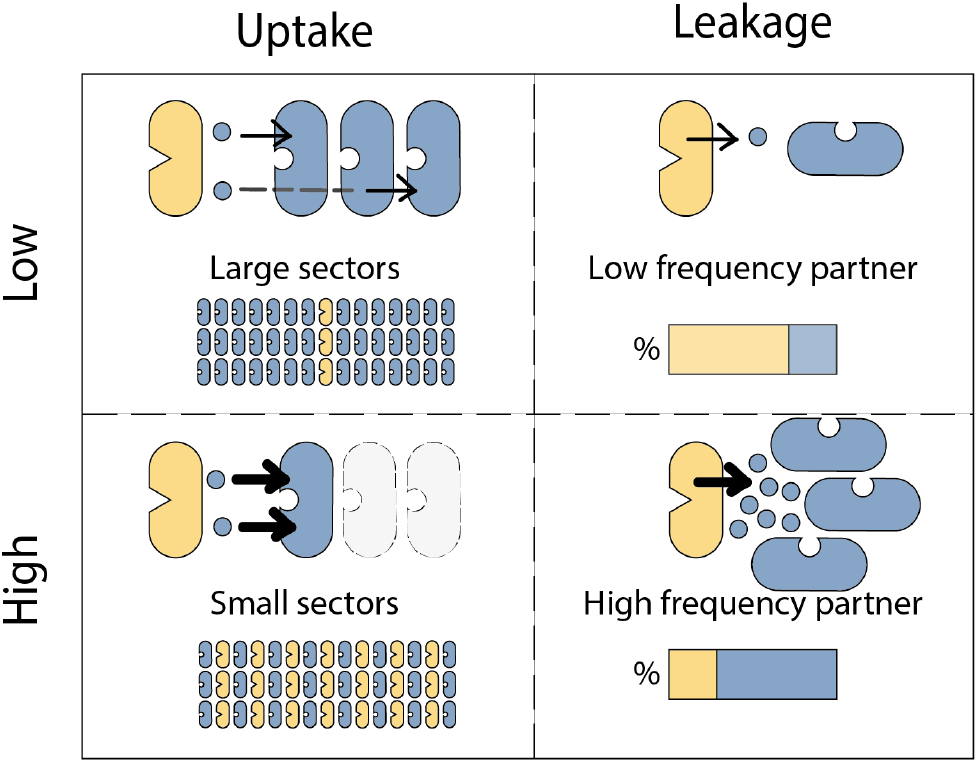
Key parameters predicted to determine community properties. Visual representation of cells exchanging amino acids (the amino acids are represented by blue circles). Uptake: Cells with a low uptake rate let a fraction of the exchanged metabolite to diffuse past them, enabling cells to grow further away from their partner and form patches of bigger size compared to cases with efficient uptake. Leakage: A low leakage rate leads to a low frequency of the partner cells and a high leakage rate to a high frequency.

Here we set out to test whether these predictions hold in a community growing in a more realistic three dimensional setting and at a bigger scale. We chose to perform range expansions on agar plates where a co-culture of proline and tryptophan *E. coli* auxotrophs can grow in three dimensions and reach up to 1 cm of radius. In addition, we used genetic engineering to test whether changing the key biophysical parameters,i.e. the uptake and/or leakage rates, indeed results in the predicted changes in spatial arrangement and/or community composition. We analysed the colonies by microscopy, followed by quantitative image analysis. We found that increasing the proline uptake rate experimentally led to smaller patches (or sectors) of the proline auxotroph. In addition, we show that an increase in the leakage rate of one or the other amino acid can change the community composition, benefiting the cells auxotrophic for the overproduced amino acid. Finally, we combined increased uptake and leakage rates and could use the model to predict the combined effect of those changes both in spatial arrangement and community composition.

## Results

### Characterising a synthetic community of two auxotrophs

We studied a synthetic community composed of two *E. coli* strains auxotrophic for amino acids (Figure 2a). The first strain is a proline auxotroph (Δ*proC*), that is unable to produce this amino acid due to a deletion of *proC* coding for an essential enzyme in the proline biosynthesis pathway. Similarly, the second strain (Δ*trpC*) cannot synthesize tryptophan, because it contains a deletion of *trpC*, which catalyses a step of the tryptophan biosynthesis pathway. While the individual strains do not grow in minimal media, the two auxotrophs can grow as a co-culture. The cells naturally leak amino acids that diffuse into the medium and can be taken up through import systems [11, 38, 39].

**Figure 2:**
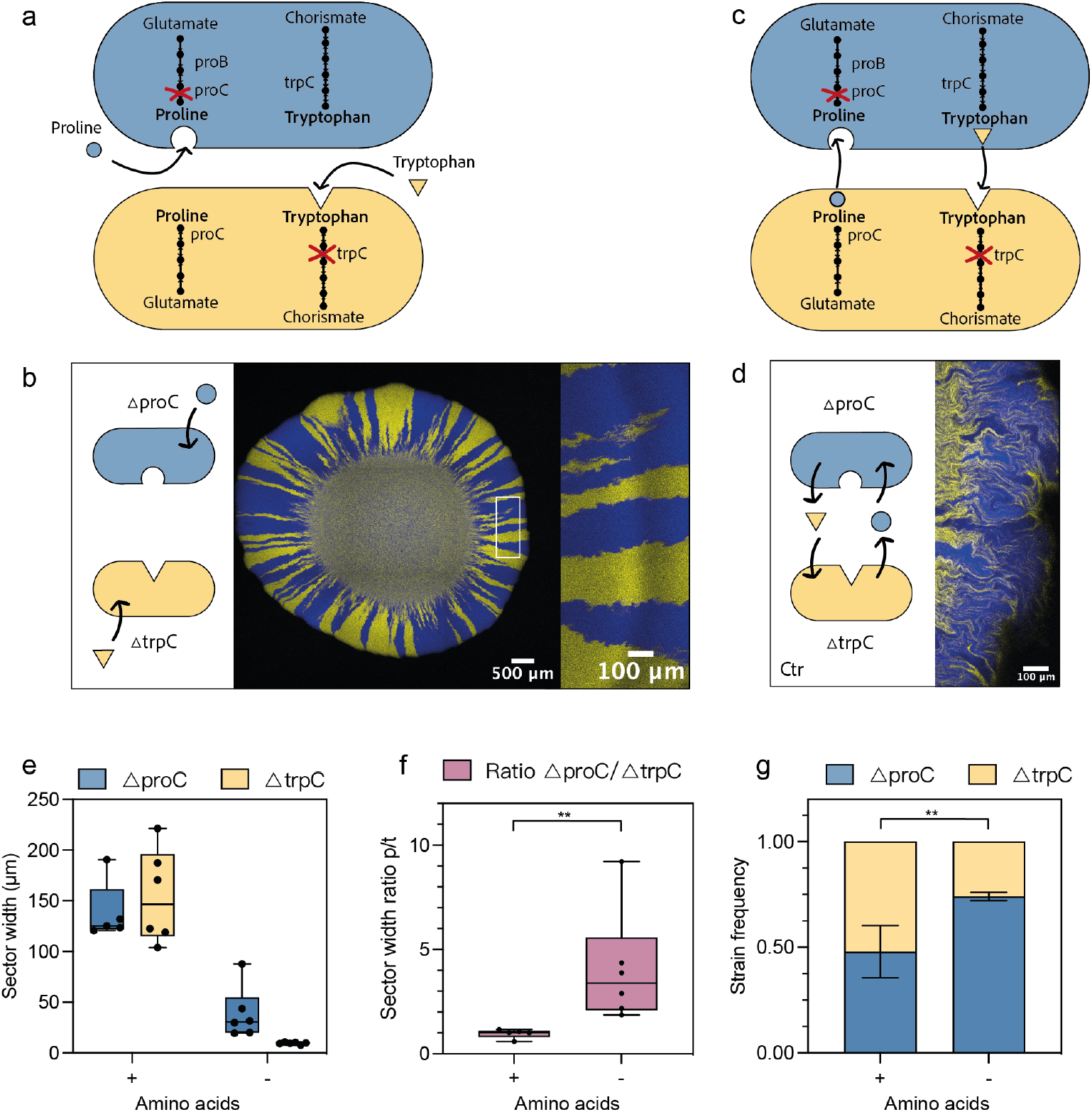
Co-culture of proline and tryptophan auxotrophs in the presence and absence of amino acids. **a** Schematic representing the relevant amino acid pathways of the strains used in this study. Knock-outs (*proC* and *trpC*) lead to proline or tryptophan auxotrophic strains, respectively. Here, proline and tryptophan were added to the medium. **b** Microscopy image of the range expansion. A representative image of one of the six biological replicates that were analysed is shown. Close-up on the edge of one of the expansions (marked with a white rectangle), at the radius where we analysed the community around the colony. **c** Schematic representation of the strains without addition of amino acids in the media, the amino acids are exchanged between the members. **d** Microscopy image of the edge of the range expansion without amino acids. **e** Quantification of the sector widths of both conditions. Box plots show the median value of the sector size of six biological replicates, error bars show min and max values. **f** Sector width ratios of proline auxotroph to tryptophan auxotroph. Box plots show the median value of six biological replicates, error bars show min and max values. Wilcoxon test, ** significant at p-value *<*0.001. **g** Quantification of the frequencies of the two communities. Six biological replicates were analysed. Student t-test, ** significant at p-value *<* 0.001.

To establish a baseline for our comparative analysis, we first characterised our co-culture in a defined minimal medium containing proline and tryptophan. In this environment, the strains do not rely on each other for growth, but compete for the resources present in the medium. To study spatial organisation and community composition during range expansion, we deposited a drop of the co-culture at a 1:1 ratio on an agar plate, and let them grow outwards for a total of five days (Figure 2b). We then imaged the range expansion by confocal microscopy. In all analyses, we only considered the edge of the expansion at the distance of 250 *µ*m from the edge of the colony where the cells grew outwards (see Figure S1 and methods for details). To quantify the spatial arrangement we measured the average sector size (Figure 2e) of each member. To facilitate the comparison of different conditions, we calculated the relative size of each strain’s sector as ratio of the *proC* strain sectors size to that of the *trpC* strain (Figure 2f). A value of 1 indicates that the sectors are of the same size. A value *>*1 indicates larger Δ*proC* than Δ*trpC* sectors, and <1 smaller Δ*proC* than Δ*trpC* sectors. Finally, we determined the relative frequency of each strain by image analysis (Figure 2g). In this initial control experiment with supplemented amino acids, both strains were equally abundant and the sectors did not differ in size. These results agree with our expectations, as the two strains have the same growth rates in this condition (Figure S2). The pattern that we observed here—well-defined sectors and a clear segregation between the members—is typical of a range expansion of two strains competing for resources [40–43].

Next, we characterized the same synthetic community in the absence of added amino acids (Figure 2c). We observed overall less growth (Figure S3) and 15-20 times smaller sectors, i.e. much higher mixing between the two strains (Figure 2d) compared to the control where amino acids were supplemented. This was expected as now the cells can only grow in close proximity to their partner. Also, the Δ*proC* sectors were wider (38*±* 18.1 *µ*m (mean*±* SE)) than the Δ*trpC* sectors (9.6 *±*0.45 *µ*m) (Figure 2e). The ratio of the sector sizes (Δ*proC* /Δ*trpC*) is 4.1*±* 1.1 (Figure 2f). Moreover, the community was asymmetrical, with a majority of Δ*proC* (74%) (Figure 2g). The observed asymmetry in sector size and community composition agrees well with the previously observed patterns of the same community grown in microfluidic devices [36]. The sector widths in the range expansions (38 and 9.6 *µ*m for Δ*proC* and Δ*trpC*, respectively) are of the same order of magnitude as the measured interaction ranges in the microfluidic chambers (12 and 3.2 *µ*m, respectively). Moreover, the sector width ratio (4.06 *±*1.1) quantitatively agrees with ratio of the previously measured interaction ranges (4.1*±* 0.42). Finally the observed community composition in the range expansions (26%*±* 0.3 of Δ*trpC*) is close to that observed in the microfluidic chambers (24.5%*±* 0.2 of Δ*trpC*) [36].

Previously, a mathematical model was developed to quantitatively predict the properties of this auxotrophic community from key biophysical parameters, primarily the uptake and leakage rates of the amino acids [37]. This model was parameterized using literature values for the uptake and diffusion rates, and we expect these rates to be the same in the range expansion setup. In contrast, leakage rates were previously fitted to the observed growth dynamics in microfluidic chambers, and we expect these rates to differ in the range expansion setup: in microfluidic chambers almost all leaked amino acids are taken up by neighboring cells, whereas on agar plates a large fraction of the molecules diffuse away into the agar. The agar thus acts as a sink, reducing effective leakage rates. To account for this, we refitted leakage rates to match the observed community composition and growth in the absence of added amino acids (Figure S4, Table S4).

### The sector width is determined by the rate of amino acid uptake

We then investigated the effect of increased amino acid uptake on the spatial arrangement and the frequency of the community members. Our model predicts that we should observe a decrease in sector width for the member which increased its amino acid uptake while the other one should stay of similar width. To test these predictions experimentally, we constructed a plasmid containing the *putP* gene, coding for a Na^+^/L-proline transporter, an active proline importer in *E. coli* [44, 45]. The overexpression of *putP* increases the proline uptake in the Δ*proC* strain. In a range expansion of this Δ*proC* + *putP* strain in co-culture with Δ*trpC* (Proline uptake community, Pu), we observed 4.4-fold thinner Δ*proC* + *putP* (blue) sectors (Figure 3c, Pu), while the Δ*trpC* (yellow) are of similar size (1.22-fold change). Consequently, the ratio of the Δ*proC* to Δ*trpC* sector widths was significantly reduced in the high proline uptake community (0.88 *±* 0.41) (Figure 3c, Pu).

**Figure 3:**
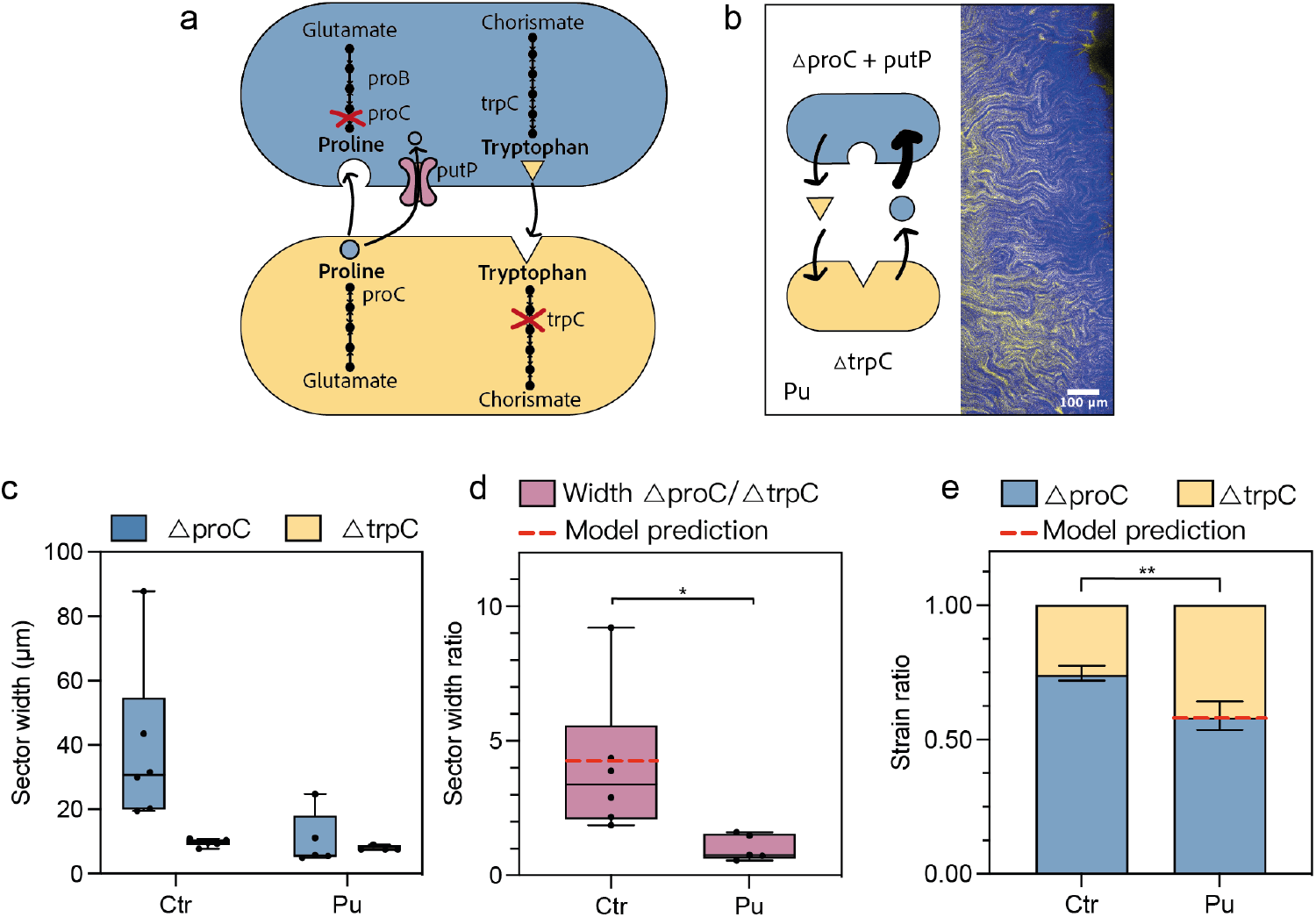
Increased proline uptake leads to smaller sectors of the proline auxotroph. Community with increased proline uptake (Pu) in the proline auxotroph. **a** Detailed view of the auxotroph community with proline importer (putP) overexpressed, leading to increased proline uptake. **b** Microscopy images of the range expansions. One representative image out of six biological replicates is shown. **c** Quantification of the sector widths in the proline uptake community (Pu) for the proline (Δ*proC* +putP) and the tryptophan (Δ*trpC*) auxotrophs. **d** Ratios of sector width of proline auxotroph to tryptophan auxotroph. The box plot shows the median value of six biological replicates, error bars show min and max values. Wilcoxon test, * significant at p-value *<*0.01. The red line shows the ratios predicted by the model. **e** Quantification of the frequencies of the two community members. Six biological replicates were analysed. Student t-test, ** significant at p-value *<* 0.001. The red line shows the frequencies predicted by the model considering the burden of putP production measured experimentally.

For the effect on the community composition, we expected that the increase in uptake rate should cause the community composition to skew further in favor of the majority type, i.e. increasing the frequency of Δ*proC*, as cells interact with fewer neighbors. However, contrary to our initial expectation, we observed a decrease in Δ*proC* frequencies (Figure 3e). We thus hypothesised that this shift is likely caused by a burden coming from the overexpression of *putP*. Indeed, the Δ*proC* + *putP* strain has a strongly decreased growth rate in liquid monoculture (80% growth rate reduction) compared to Δ*proC* (Figure S2). We incorporated the experimentally measured cost of *putP* overexpression in the model. Moreover, we estimated the increase in proline uptake rates by fitting the model to the observed sector width ratio. With this addition, the predictions agree with the observed decrease of Δ*proC* + *putP* frequency (Figure 3e, Pu). In summary, by overexpressing *putP* we could confirm the importance of the uptake rates in shaping the spatial arrangement of the synthetic community.

### Amino acid overproduction leads to a change in community composition

Secondly, we studied the effect of amino acid availability on the community, as this parameter was predicted to have an impact on the community composition [37, 46]. We constructed two strains that overproduced the amino acid required by their partner strain. We engineered a proline auxotroph overproducing tryptophan and a tryptophan auxotroph overproducing proline (Figure 4a). The tryptophan overproduction was obtained by deleting *trpR*, a repressor of the tryptophan biosynthesis pathway [47]. The proline overproducer contains a point mutation (A319G) in the *proB* gene as this mutation (designated as *proB74*) was reported to lead to overproduction of proline in *E. coli* due to a loss of allosteric regulation [48]. Figure 4b shows the pattern of the expansions of three overproducer communities: 1) T_p_: tryptophan overproducer (Δ*proC* ΔtrpR) with Δ*trpC*, 2) Pp : proline overproducer (Δ*trpC proB74*) with Δ*proC* and 3) Tp Pp : both tryptophan (Δ*proC* ΔtrpR) and proline overproducers (Δ*trpC proB74*). Interestingly, we observed much larger population size for the communities with the proline overproducer (Pp and Tp Pp) (Figure S3). Consequently, the absolute values of the sector sizes were much higher for those communities (Figure 4c). Still, in all cases, the sectors of the proline auxotroph strains were wider than those of the tryptophan auxotroph strains. Indeed, as predicted by the model, there was no important change in the ratio of the sector sizes (Figure 4d) for any of the communities (p-values between 0.177-0.015).

**Figure 4:**
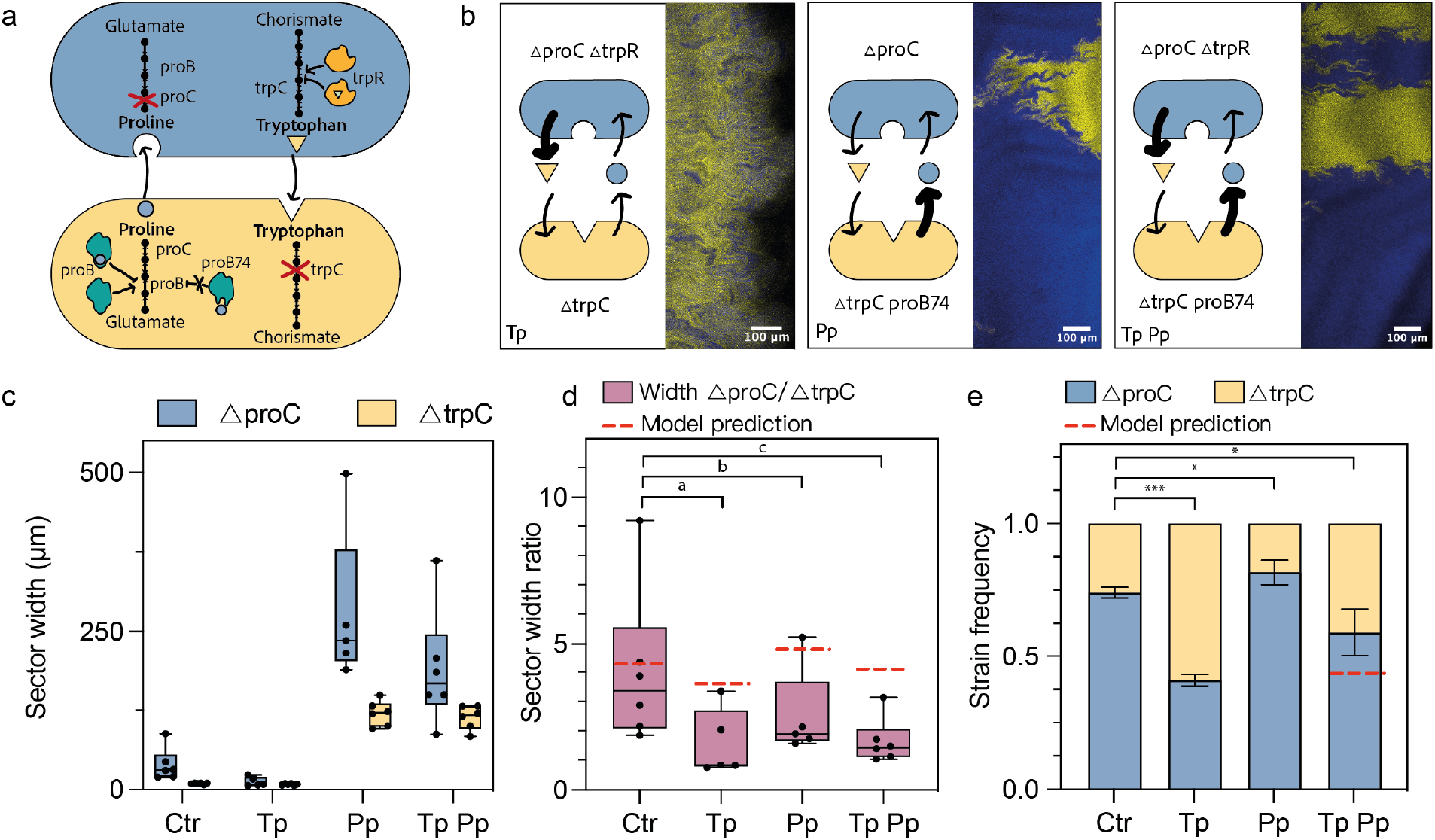
Amino acid overproduction leads to a change of community composition. Comparison between the initial community (Ctr) and communities with increased production of tryptophan (Tp), proline (Pp) or both (TpPp). **a** Detailed view of the metabolic pathways and modifications in the communities used in this assay. *TrpR* regulates the tryptophan pathway, a deletion of its gene leads to overproduction of tryptophan. *ProB* regulates the proline pathway, and the mutation *proB74* leads to proline overproduction. **b** Microscopy images of the range expansions. One representative image out of six biological replicates is shown. **c** Quantification of sector widths of each community for six biological replicates per condition, boxplot shows median and error bars show min and max values. **d** Ratio of sector widths for each community. Wilcoxon test (p-values: a=0.05195, b=0.1775, c=0.01515). The red lines show the ratios predicted by the model. **e** Quantification of community composition for six biological replicates. Student t-test, * significant at p-value *<*0.01 and *** significant at p-value *<*0.0001. The red line shows the frequency predicted by the model.

Next, we analysed the community composition that was predicted to change in response to the increased leakage caused by amino acid overproduction. Indeed, we observed an increase in the frequency of the strain benefiting from the overproduced amino acid (Figure 4e): Δ*trpC* increased in frequency when paired with the tryptophan overproducer in the Tp community and Δ*proC* showed a higher frequency when paired with a proline overproducer in the Pp community. We used the observed community composition in the Tp and Pp communities to estimate the increase in tryptophan and proline leakage rates, respectively, and used these values to predict the expected composition in the Tp Pp community. The predicted frequency of 50% Δ*proC* is close to the observed value of 59% Δ*proC* and falls in between the two individual overproducer communities. In conclusion, by overproducing the amino acids we demonstrated the importance of the leakage rates in determining the composition of the synthetic community.

### The effects of uptake and overproduction are cumulative

Next, we combined the changes in uptake and leakage rates. Our expectation was that we would change both sector widths and community composition. In particular, an increased proline uptake should lead to smaller Δ*proC* sectors and a higher production of one amino acid should increase the frequency of the partner strain. We thus tested all the combinations with the strains that we had constructed. Namely, tryptophan overproducer and increased proline uptake (Tp Pu), proline overproducer and increased proline uptake (Pp Pu) as well as both tryptophan and proline overproduction with an increased proline uptake (Tp Pp Pu) (Figure 5a,b). As we had observed previously, communities containing the Pp grew more than the control community (Figure S3), leading to overall bigger sectors (Figure 5c). In agreement with the expectation for Pu, all communities had reduced sector width ratios compared to the control (Figure 5d). For the community composition (Figure 5e), we observed a significant increase of the Δ*trpC* frequency in the Tp Pu community due to the increased production of tryptophan as predicted by our model. For the Pp Pu community we measured a slight decrease in the Δ*proC* frequency rather than an increase due to the increased proline production. However, taking into account the burden of overexpressing *putP*, the result agrees well with the model prediction. Finally, for the community with all three modifications, Tp Pp Pu, we found that the proline auxotroph decreased in frequency to 58% (Figure 5d). Although the model correctly predicted the direction of change, it overestimated the magnitude of the change, likely because amino acid concentrations are no longer growth limiting for all cells in this in this community (see discussion). To summarise, we have shown that the combination of uptake and leakage rates is cumulative and changes the community arrangement and composition accordingly.

**Figure 5:**
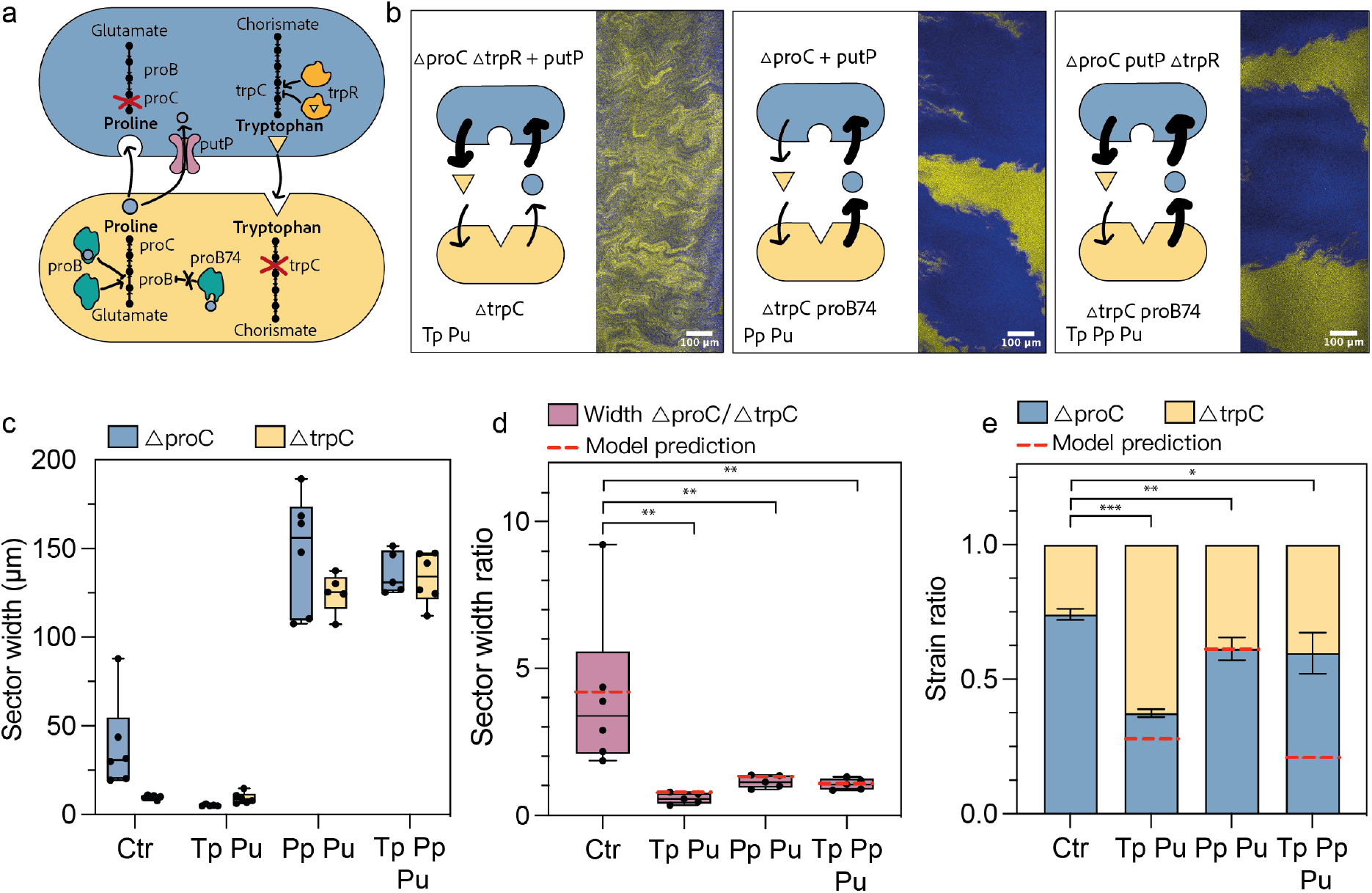
Combined effect of increased amino acid uptake and overproduction. Comparison between initial community (Ctr) and communities with tryptophan overproduction and increased proline uptake (Tp Pu), proline overproduction and increased proline uptake (Pp Pu), and both tryptophan and proline overproduction and increased proline uptake (Tp Pp Pu). **a** Detailed view of the metabolic pathways and modifications in the communities used in this assay. **b** Microscopy images of the range expansions. One representative image out of six biological replicates is shown. **c** Quantification of sector widths of each community. Box plots show the median of six biological replicates, error bars show min and max values. **d** Ratio of sector widths for each community. Wilcoxon test, ** significant at p-value *<*0.001. The red lines show the ratios predicted by the model. **e** Quantification of community composition. Student t-test, * significant at p-value *<*0.01, ** significant at p-value *<*0.001, *** significant at p-value *<* 0.0001. The red lines show the frequencies predicted by the model.

## Discussion

In this study, we described how two main parameters—amino acid uptake and leakage rates—determine a community’s spatial arrangement and composition (Figure 1). The use of a simple synthetic community (Figure 2), combined with genetic engineering of the strains, allowed us to modify these two variables independently. In agreement with a previously proposed model [37], we experimentally demonstrated that increased amino acid uptake leads to smaller sectors, and thus a more mixed community, as the cells have to be close to each other to benefit from the shared molecule (Figure 3). An increased amino acid leakage by one cell type leads to a higher frequency of its partner as the amino acid it is lacking is present at a higher concentration (Figure 4). Changes of the two parameters can be combined to control spatial arrangement and composition at the same time (Figure 5).

While overall our model predicted very well the changes that we observed experimentally, in a few cases, the predictions deviated from the experimental results (Figure S4). Specifically, the model could qualitatively predict the observed direction of change in equilibrium frequencies and sector width ratios for nearly all communities. For most communities there was also a good quantitative agreement between the model predictions and the experimental observations, however predictions were much less accurate for the communities that contain both proline and tryptophan overproducers (Tp Pp and Tp Pp Pu). This can be explained as the model assumes that amino acids concentrations are always very low, i.e. well below the saturation constants of both the uptake and growth functions. This assumption is likely violated in communities that contain both overproducers and this causes the model to fail in two ways: First, the model overestimates the uptake rates in the Pp and Tp communities, and this leads to an underestimate of the range over which cells can interact. As a result the model cannot recapitulate the large increase in Δ*proC* and Δ*trpC* sector widths in the communities containing Pp (Figure 4c). This also explains why we fail to accurately predict the sector width ratio for the Tp Pp community. Second, the model underestimates the growth rate of cells when amino acid concentrations reach growth saturating levels, as it assumes that the growth rates of cells increase linearly with the local frequency of the partner type. Here again, this assumption is probably not valid in communities containing both overproducers and therefore the model did not estimate the strain frequencies correctly for the Tp Pp Pu community (Figure 5e). Although our model could be adapted to include saturating uptake and growth functions, this would complicate the mathematical analysis, and we thus leave such an extension for future work. Finally, the range expansion setup used in the experiments, breaches another important assumption of the model: namely that all of space is occupied by cells. The model was originally developed for densely packed populations such as those found in 2D microfluidic growth chambers or in the cores of 3D biofilms. In these systems metabolites can only diffuse in the narrow spaces between cells. However, on agar plates metabolites can also diffuse through the agar leading to an increased range of diffusion. This is a second reason why our model generally underestimates the observed sector widths. Moreover, the agar acts as a sink for the amino acids leaked from the cells. This means that the effective leakage rate is different to the one previously calculated in microfluidic chambers. To solve this, we have refitted the leakage rates on our control communities (Ctr, Tp and Pp). In summary, although there were a few exceptions where the model was less accurate, which we can explain, we experimentally demonstrated that leakage and uptake rates influence community arrangement and composition in a more realistic 3D setting and on a larger scale than the microfluidic setup used to develop the theory.

In this study, we focused on the edge of the range expansion, where the community grows and needs to exchange amino acids. At the edge, most cells grow outwards, where the nutrients are abundant and the space is free. This resembles the situation in a microfluidic chamber where the nutrients are only accessible from one side of the chamber and the cells typically grow in one direction only [49]. In contrast, in the centre of a range expansion, the cells grow in all directions on the agar. However, they quickly merge with other microcolonies, which hinder their growth as they compete for space and nutrients. Typically, at the densities we worked with, the center was filled after 24 to 48 hours while the edge grew for up to five days. For future work, it would be interesting to study if the same rules also apply when growth occurs in all directions, as in the center of the range expansion experiments.

Do uptake and leakage rates also play a similar role for other auxotrophic communities? Preliminary results indicate that our observation might also hold true for other synthetic communities of amino acid auxotrophs. We tested a histidine auxotroph (ΔhisD) and a methionine auxotroph (ΔmetB) in combination with Δ*proC*. Again, we used *proB74* for proline overproduction and also built a histidine overproducer by deleting *hisL*, which had been shown to result in histidine overproduction by releasing the negative regulation of the pathway [47]. We were able to show that the frequency of the community members was changing according to the amino acid production level when proline or histidine were overproduced (Figure S5). Similarly to proline and tryptophan, we have modified these auxotroph strains to overexpress specific importers for histidine and methionine (hisJPQM and metNIQ)[50, 51]. While we detected differences in patterns, we could not quantify them with our current imaging and analysis pipeline, as the sectors are extremely small. We thus have some evidence that the same rules apply for other communities of amino acid auxotrophs, however, further research is required to see how general the uptake and leakage rates of exchanged metabolites determine spatial arrangement and composition of microbial communities.

While, here, we studied synthetic communities in a controlled environment, this simple model system allowed us to understand principles that we believe are also relevant for natural situations and that would have been difficult to unravel in a more complex setting. Indeed, range expansions experiments like ours have been previously used to decipher ecological processes taking place in diverse microbial communities. These studies typically focused on known interactions within a community and described their effect on factors such as pattern formation, expansion radius or frequency of partners [30, 32, 33, 41, 52, 53]. Additionally, the initial conditions of the range expansion have been modified such as initial ratio or density of cells, resources availability or antibiotic stress [27, 40, 54–56xb
]. Finally, genetic modifications have been applied to community members showing the effect of a specific feature of the cells on the final pattern, such as motility [42]. In our study, we show the effect of the interactions between two strains via the exchange of amino acids—metabolites whose exchange is also relevant in a variety of natural communities ([11, 16–19]. Moreover, we have shown that modifying the strength and range of interactions by genetic modifications is a powerful approach to understand how they influence the microbial community. The parameters that we chose to modify, uptake and leakage rates, are also subject to change during evolution. For example, a recent laboratory evolution experiment with a microbial community of *E. coli* and *S. cerevisiae* amino acid auxotrophs, selected mutations which led to increased uptake and production rates of the exchanged amino acids [57].

In addition to improving our understanding of how molecular interactions between community members determine community-level properties, the results that we describe here are also relevant for applications where the tight control of community composition is desirable. For example, it is well established that distributing the bioproduction across a synthetic consortia can increase the yield compared to production in a single strain [58–60]. However, maintaining the stability of these engineered communities remains a challenge. Among other strategies, cross-feeding between community members has been used to establish stable communities [61, 62]. Similarly to the results presented here, we have observed that we can also control the frequency of members of our synthetic community in liquid cultures by changing leakage rates (Figure S6). Modifying leakage and uptake rates might therefore be a promising approach to control the composition and spatial arrangement of synthetic co-cultures for bioproduction [63–66], but also for other applications such as food production [67], distributed biocomputing [68] or engineered living materials [69].

## Material and methods

### Bacterial strains

All the experiments performed in this study were done using strains derived from *E. coli* MG1655 constitutively expressing sfGFP or mCherry and carrying the deletions Δ*trpC* and Δ*proC*, respectively (kindly provided by A. Dal Co, [36, 70]). Details of all strains are listed in Supplementary Table 1. All further gene deletions were performed with the prophage-lambda red recombination system as described previously [71]. The linear donor DNA fragment for the homologous recombination carried an antibiotic resistance gene and FRT sites. It was amplified from pKD3 or pKD4 by PCR (2x Phanta Max Master Mix, Vazyme) introducing extensions that are homologous to regions adjacent to the gene to be inactivated. After recombination, the antibiotic resistance gene was eliminated following the lamda red protocol. The primers used are listed in the Supplementary table 3.

For the Tp strain we introduced a *trpR* deletion. The linear donor DNA fragment was amplified from pKD3 using primers prEP167 and prEP168. The Pp strain was generated by mutating a single nucleotide (A319G) in the *proB* gene [48]. To introduce this mutation, first, the *proBA* operon was deleted from the genome by recombination of a donor fragment amplified from pKD3 using the primers prEP197 and prEP198. We removed the antibiotic cassette following the lambda red protocol. Then we built pEP27, a plasmid derived from pKD4, harbouring the FRT sites, the kanamycin resistance cassette and the *proBA* operon, including the single nucleotide mutation on proB. The plasmid backbone was obtained by amplifying pKD4 by PCR using the primers prEP177 and prEP200. *proBA* was amplified from the genome of MG1655 in two pieces: *proB* until the mutation, and *proB* from the mutation to *proA*, using prEP201 and prEP185, and prEP203 and prEP202. The 3 fragments were assembled using Gibson assembly (NEBuilder HiFi DNA Assembly Master Mix, NEB). To re-introduce the proBA operon (now with A319G) mutation on *proB*, we amplified the sequence FRT-Km-FRT-proBA(A319G) with the primers prEP205 and prEP250 and then used again the lambda red protocol [71].

The histidine auxotrophic strain was obtained by deletion of *hisD* using prEP207 and prEP208 (lambda red recombination amplifying the fragment from pKD3) and the histidine overproducer (Hp) was obtained by knocking-out the *hisL* gene, using prEP227 and prEP228 (lambda red recombnination, amplifying the fragment from pKD3). The methionine auxotrophic strain was obtained by deletion of *metB* using prEP211 and prEP212 amplifying from pDK3.

### Plasmids

All strains in this study harbour either a plasmid with the anhydrotetracycline (aTc) inducible expression of the *putP* gene (pEP17) or an identical control plasmid (pEP28) except that it did not contain *putP*. These plasmids are presented in Supplementary table 2. pEP17 was built using the backbone of pDSG360, which was a kind gift from Ingmar Riedel-Kruse (Addgene plasmid #115601) [72]. The primers prEP111 and prEP112 were used to amplify by PCR the relevant parts of pDSG360. Primers prEP113 and prEP114 were used to PCR amplify *putP* from the genome of MG1655. The plasmid was assembled by Gibson Assembly (NEBuilder HiFi DNA Assembly Master Mix, NEB). The control plasmid pEP28 was built from pEP17, by amplifying it without the *putP* gene with primers prEP206 and prEP111 and assembled with Gibson assembly (NEBuilder HiFi DNA Assembly Master Mix, NEB).

### Range expansions experiments

Range expansions were performed in six replicates. For each replicate, starter cultures were obtained by inoculating a single bacterial colony in 2 mL of lysogeny broth (LB) supplemented with chloramphenicol (50 µg/mL) and overnight incubation at 37 °C with orbital shaking (200 rpm). The next morning, each culture was refreshed 1:100 (20 µL of starter culture in 2 mL of fresh LB medium) and grown for 4 h in identical incubation conditions. Cells were then washed twice with an equal volume of PBS (i.e. 500 µL of culture were washed with 500 µL of PBS) and mixed at a 1:1 ratio at an optical density (OD_600_) of 2. Agar plates were prepared 24 h in advance to ensure reproducible humidity of the plates. From the co-culture, 1.5 µL were gently pipetted on M9 minimal medium plates (1.5% agar, 47.76 mM Na2HPO4, 22.04 mM KH2PO4, 8.56 mM NaCl, 18.69 mM NH4Cl, 1 mM MgSO4, 0.1 mM CaCl2 and 10 mM glucose) supplemented with chloramphenicol (50 µg/mL) and aTc (50 ng/mL), where needed we added 50mg/L of proline and 20mg/L of tryptophan. Plates were placed in a 37 °C incubator for 5 days. The range expansions were imaged with a Zeiss LSM 900 confocal microscope. sfGFP and mCherry were excited at 488 nm (0.8% intensity) and 561 nm (0.65% intensity) using a 45 µm pinhole, respectively. A 5x objective was used at 1.5x zoom, 15 Z stacks were taken for each image as well as 4 or 9 tiles depending on the size of the picture. The stitching was automatically carried out using the Zen Blue software.

### Image analysis

We measured the sector size and strain ratios at 11 randomly chosen positions for 6 different range expansions for each condition. We averaged the 11 measurements to have one value for each biological replicate. To do this, using Fiji [73], we first projected all 15 z-stacks to one image (Z-projection, Max intensity). Then, we analyzed the image using a custom-made pipeline. First, the centre and the boundary of the colony were found with the *regionprops* function of MATLAB. Based on the slightly elliptical shape of the colonies, the properties (such as the separation length or the molar ratio) were calculated on concentric ellipses, while the radius of the colony (*R*) was defined as the average of the major and minor radius of the ellipse. The volume of the initial droplet was the same in every experiment, so it was assumed that the radius of the inner ring was the same in every case as well: *R*_*c*_ = 1.89·10^3^ *µm*. The *growth length* of the colony was defined as the difference between the radius of the colony and the radius of the inner ring: *l*_*growth*_ = *R*−*R*_*c*_. Figure S7 illustrates the process of the image analysis: the magenta curve in **a** shows the edge of the colony. Due to the stochastic nature of colony growth, this boundary is often not smooth, furthermore, the properties are changing rapidly around the edge, so the phenomenological properties of the colonies were calculated on a concentric ellipse with a 250 *µm* distance from the edge of the colony (blue curve): *R*_*calc*_ = *R*− 250 *µm*. The autocorrelation function was calculated around the concentric ellipses. The *sector width* (*s*) was defined as the shortest distance when the value of the autocorrelation function goes below a given threshold (0.2). This threshold was manually chosen, but we confirmed that all results are robust to its precise value. In the case of the community of the strains without auxotrophies, strong separation was observed, resulting in a continuous increase in sector size and the separation length (Figure S7b). When the separation was weaker, the noise in the separation length function was bigger, and the analysis of multiple images and colonies was necessary to get reliable results. In the calculation of the average separation length for the colonies, 11 data points were averaged around the previously determined radius, *R*_*calc*_ (blue, dashed, vertical line). The ratio of the red cells (*x*_*red*_) (Δ*proC*) was calculated around the concentric ellipse as well. In this case, this ratio was determined based on the sum of the red (*R*) and green (*G*) fluorescent intensities along the ellipse: *x*_*red*_ = *R/*(*R* + *G*) (Figure S7c). The average *red ratio* for a colony was calculated from 11 points around *R*_*calc*_. In some cases, our pipeline resulted in clear outliers most likley due to noise. To reduce their impact, we employed a robust fitting method using bisquare weights with MATLAB’s “fit” function [74]. A constant function was fitted, points further from the fitted curve receive ligher weights. The points significantly distant (what could not be expected by random chance), receive zero weight. We identified the points with zero bisquare weight as outliers and excluded them from our data.

### Model

The full model and its derivation were previously described in reference [37]; here we only summarize the most important assumptions and results.

#### Model assumptions

The model is based on the following main assumptions:

1. All cells take up amino acids with a constant uptake rate *u*. Amino acid concentrations are assumed to be low enough such that saturation of importers can be ignored.
2. Growth is limited by the amino acid for which a cell is auxotrophic. The growth rate *µ* follows Monod kinetics: 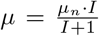, where *I* is the internal concentration measured in units of the Monod constant, and *µ*_*n*_ the maximum growth rate of the auxotroph in the presence of amino acids.
3. Producer cells maintain a constant internal concentration of *I*_*C*_.
4. Cells externalize amino acids into the environment through passive diffusion across the cell membrane with leakage rate *l*.
5. Cells are homogeneously distributed through all of space. Molecules can only diffuse through the space between cells, and the effective diffusion rate is given by 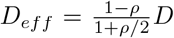, where *ρ* is the cell density and *D* the diffusion constant in solution [75].

These assumptions were based on the prevailing conditions in densely packed communities of auxotrophic cells, such as those found in 2D microfluidic growth chambers or in the core of 3D biofilms. However, we expect most of these assumptions to hold for the range expansion setup as well. One exception, is the 5th assumption: in the range expansions setup metabolites can also diffuse into the agar, thus reducing effective leakage rate, and increasing the effective diffusion range (see Discussion). Moreover, the first assumption, that uptake is non-saturated, could potentially be violated in the overproducing communities (see Discussion).

#### Predicting sector width ratio

Based on these assumption, the range over which amino-acids are exchanged can be calculated analytically. We previously showed that this interaction range, *R*, is given by:

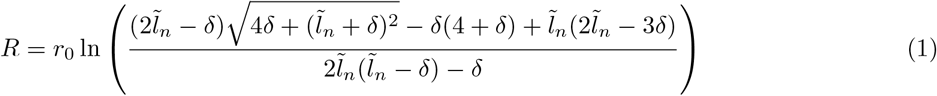

Where 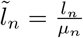 is an auxotroph’s leakage rate relative to its growth rate and:

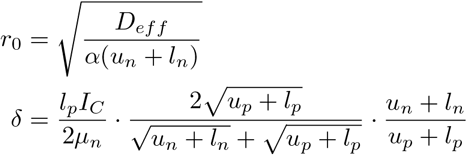

where *u*_*p*_ and *l*_*p*_ are the uptake and leakage rates in the amino acid producing cells, *u*_*n*_ and *l*_*n*_ the rates in the non-producing auxotrophs, and *α* = *ρ/*(1− *ρ*) is the volume ratio of the intra-to extracellular space. Equation 1 above is equivalent to Eq. 8 in reference [37], however, we did not use the simplifying assumption that leakage rates are low, as this is not necessarily the case for the communities containing overproducers.

The interaction range does not directly equal the sector width in the range expansion experiments, however we do expect them to be proportional to each other. An increase in interaction range allows auxotrophs to grow in regions further from producer cells and should thus lead to larger sectors. To compare the model to the data we thus compare the predicted ratio of the Δ*proC* to Δ*trpC* interaction ranges (i.e. 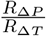)to the observed ratio of sector widths.

#### Predicting community composition

Moreover, we previously showed that the equilibrium frequency of the community is given by:

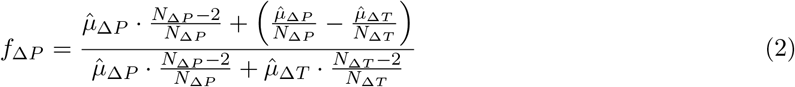

where *f*_Δ*P*_ is the final frequency of the proline auxotrophs. 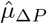 and 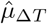 in equation 2 are the maximum growth rates which the proline and tryptophan auxotrophs can reach when they are fully surrounded by amino acid producing cells. We previously showed that this maximum growth rate is given by:

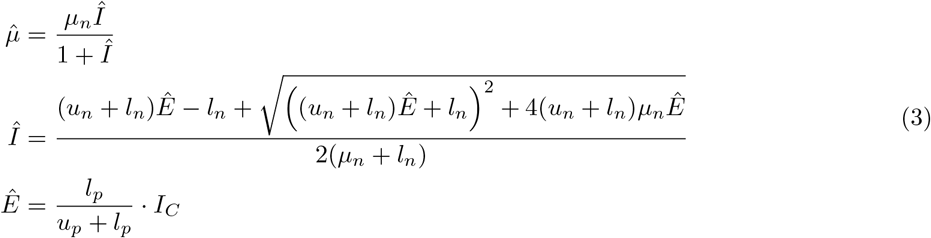

here Ê corresponds to the external concentration of the leaked amino acid in a region of space that is fully occupied by producer cells, and Î is the corresponding internal concentration of that amino acid in a isolated auxotrophic cell in that a region (both measured in units of the Monod constant). Equation 3 is equivalent to equation 7 in reference [37], however, we did not use the simplifying assumption that leakage rates are low.

Moreover, *N*_Δ*P*_ and *N*_Δ*T*_ in equation 2 are the number of cells with which the proline and tryptophan auxotrophs interact, respectively. This can be calculated from the interaction range, using a simple geometric argument. We assume cells are cylinders with spherical caps with a total length *L* and diameter *W*, and that they interact with all cells that are within distance *R* from their cell surface (in three dimensions). The number of interaction partners is then given by:

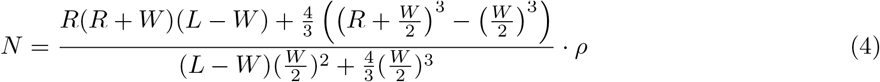

The composition of the community primarily depends on the leakage rates. Specifically, in communities where both strains have the same uptake and leakage rates (i.e. *l*_*p*_ = *l*_*n*_ and *u*_*p*_ = *u*_*n*_) the leakage rates fully determine which community member is more abundant: namely the one that depends on the amino acid that has the highest leakage flux (*l*_*p*_*I*_*C*_). However, the exact frequency of the dominant community member is influenced by the uptake rate: Higher uptake rates lead to more local interactions, skewing the composition further in favor of the dominant community member. If strains differ in their uptake and leakage rates, the community composition depends on the exact values of these rates. Generally leakage rates still have the strongest effect, however increasing the uptake rate in the auxotroph, relative to that of the producer (i.e. *u*_*n*_ *> u*_*p*_) will increase its frequency to some extent.

In deriving equation 2 we made two additional assumption: 1) cells interact equally with all other cells that are within the interaction range, and 2) leaked amino acid concentration are low enough such that auxotroph growth rate always increase linearly with the frequency of producer cells within the interaction neighborhood, without saturating. This second assumption could be violated in the case of overproducing strains.

#### Model parameterization

Model parameters are shown in Supplementary table 4. Most parameters where obtained from literature, or from previous estimates from single-cell imaging experiments. However, no accurate estimates could be obtained for the leakage rates of proline and tryptophan nor for the proline uptake rate of the *putP* mutant, so these were estimated by fitting the model to the data. The leakage rates of proline and tryptophan in the Δ*trpC* and Δ*proC* strains, respectively, where simultaneously fitted by minimizing the combined normalized Euclidean distance between the predicted and observed community composition and relative growth rate of the Ctr community. Specifically, we minimized 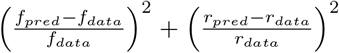, where *f*_*pred*_ and *f*_*data*_ are the predicted and mean observed frequency of Δ*trpC* in the Ctr community and where *r*_*pred*_ and *r*_*data*_ are the predicted and mean observed values of the final colony expansion range of the Ctr community in the absence of amino acids relative to that in the presence of amino acids. The leakage rates of proline in Δ*trpC proB74* and of tryptophan in Δ*proC* ΔtrpR were fitted by minimizing the Euclidean distance between the predicted and observed community composition in the Pp and Tp communities, respectively. Finally, the uptake rate in the Δ*proC putP* was fitted by minimizing the Euclidean distance between the predicted ratio of the Δ*proC* to Δ*trpC* interaction range and the mean observed ratio of the Δ*proC* to Δ*trpC* sector widths, in the Pu community. The community composition and sector width ratio are independent quantities that are primarily set by different biophysical parameters (leakage and uptake respectively). For communities on which we fitted the equilibrium composition, we can thus still predict the sector width ratio and vice-versa.

## Acknowledgements

We thank Alma Dal Co for her advice from the beginning of this project, her mentoring, and her communicative fascination for these “blue and yellow guys”. We dedicate this article to her memory. We thank Sara Mitri, Oliver Meacock, Audam Chhun and I ç vara Aor for the helpful scientific discussions and comments on the manuscript.

This work was funded by the University of Lausanne and the NCCR Microbiomes (National Centre of Competence in Research), funded by the Swiss National Science Foundation (grant no. 51NF40 180575). S.v.V was additionally supported by an Ambizione grant (grant no. PZ00P3 202186) from the Swiss National Science Foundation and by the University of Basel.

## Author contributions

E.P and Y.S designed the experimental research. E.P and T.S performed the experiments. G.H developed the image analysis method. E.P and G.H analyzed the data. S.vV designed the computational part and performed the mathematical modeling. E.P and Y.S wrote the paper with input from S.vV and G.H. All authors revised the manuscript and approved the final version of the manuscript.

## Data availability

The Python code used for model fitting and predictions is available on Zenodo: https://doi.org/10.5281/zenodo.12773415

Plasmids will be available at Addgene.

