## Supplementary Information for "Engineering microbial consortia: uptake and leakage rate differentially shape community arrangement and composition"

**Supplementary Figures**

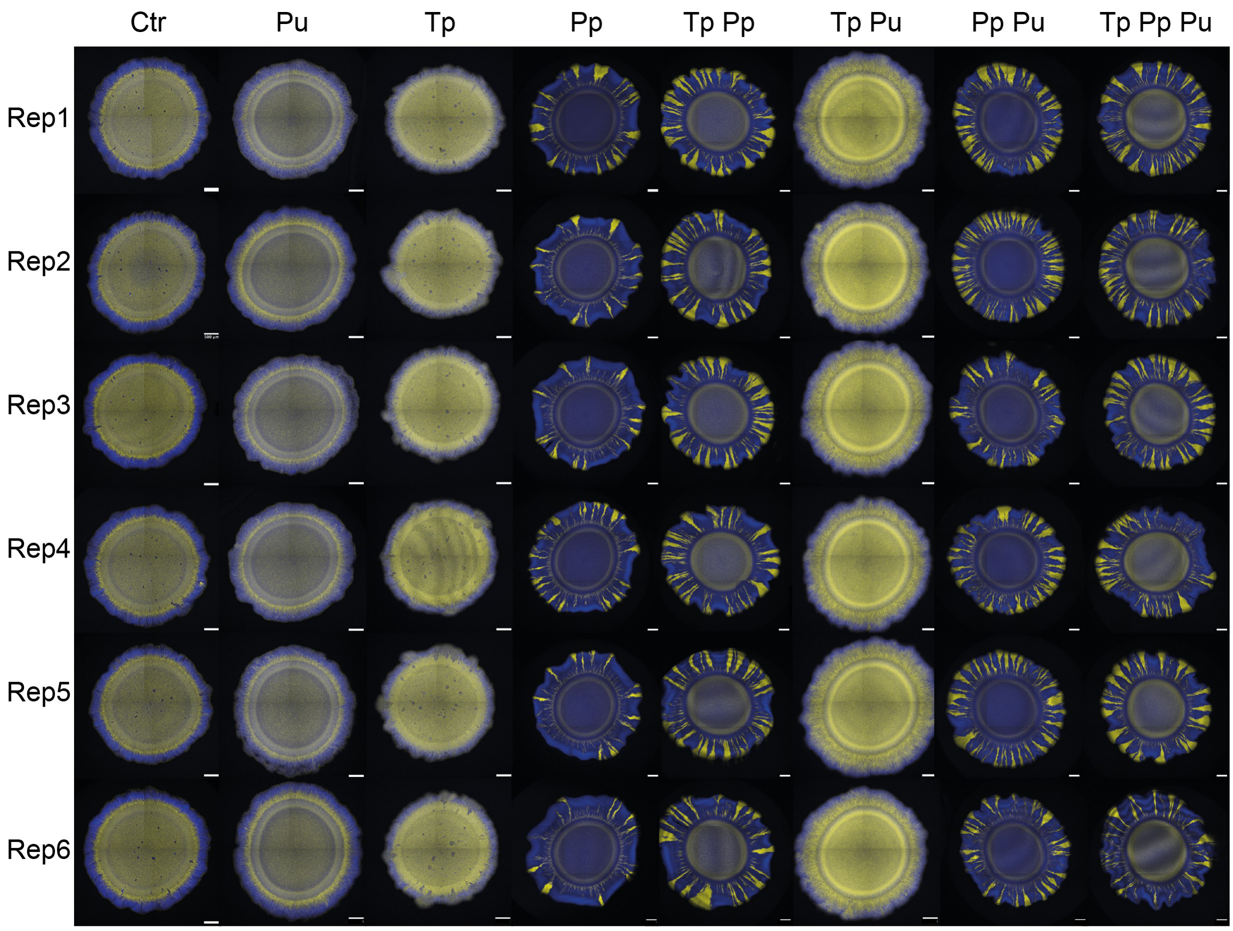


Figure S1: **Full images of the range expansions.** Full pictures of all 6 replicates for all communities analyzed in this study. Scale bars = 500$\mu$m

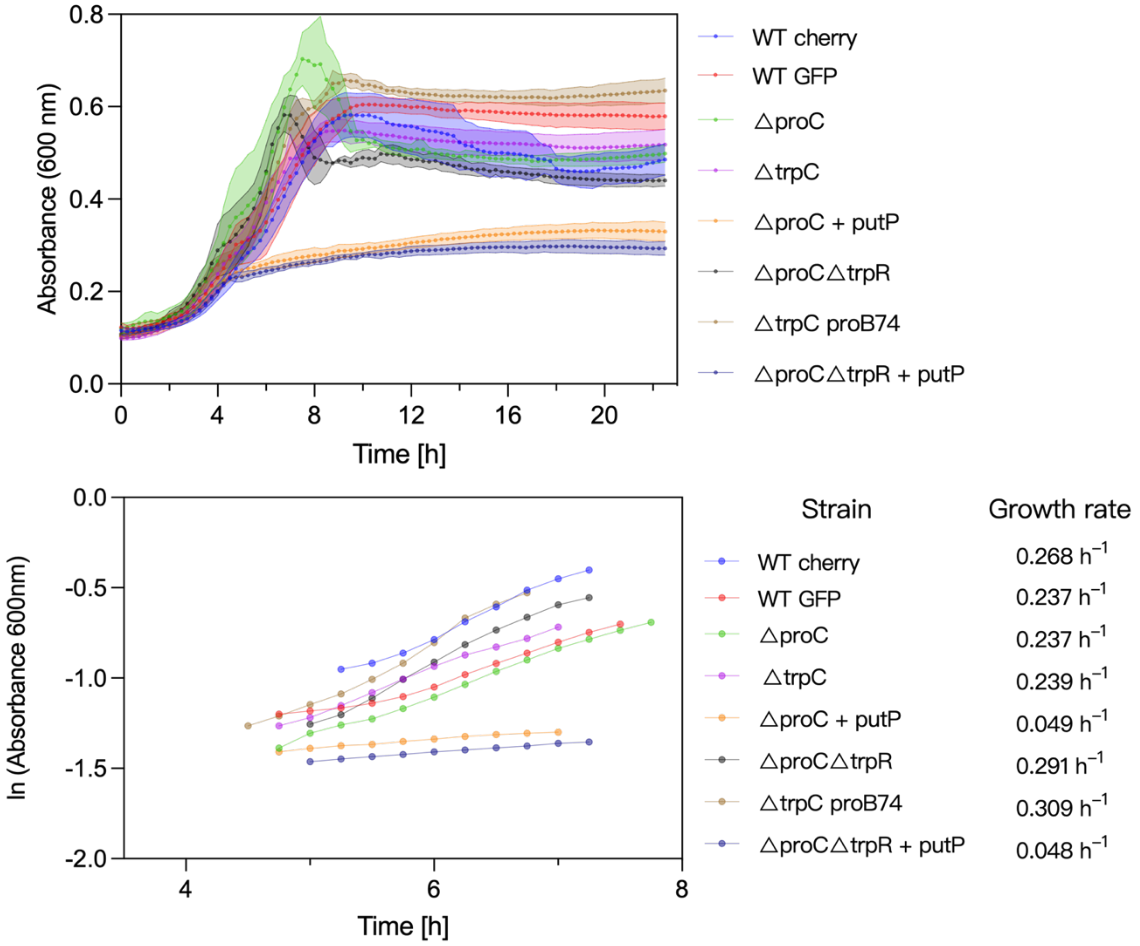


Figure S2: **Growth curves of all strain used in this study as monocultures.**
The strains were grown in M9 liquid media supplemented with proline and tryptophan (50 mg/mL and 20 mg/mL). **a** Absorbance at 600 nm measured over 22 h with 15 min between time points. Mean and standard deviation for three biological replicates for each strain. **b** Growth rates. First, we plotted the natural logarithm of the absorbance data points (ln (Absorbance 600 nm)), then isolated the exponential growth phase of the curves by selecting the part of the curves that fitted the linear trendline with a $R^2$ higher than 0.9 and took the slope of this curve as growth rate (right column).

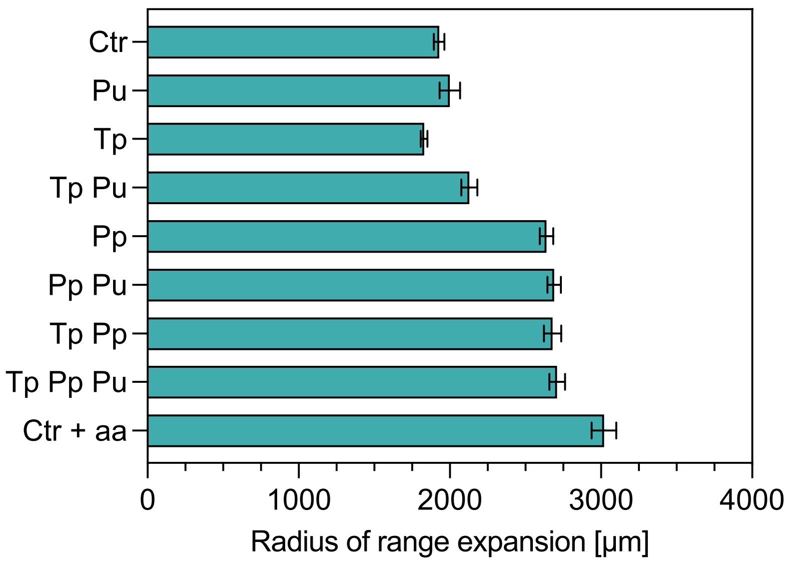


Figure S3: **Radius of the range expansions**. Growth ranges in $\mu$m for all the different communities presented in this study. Mean and standard deviation of 6 biological replicates.

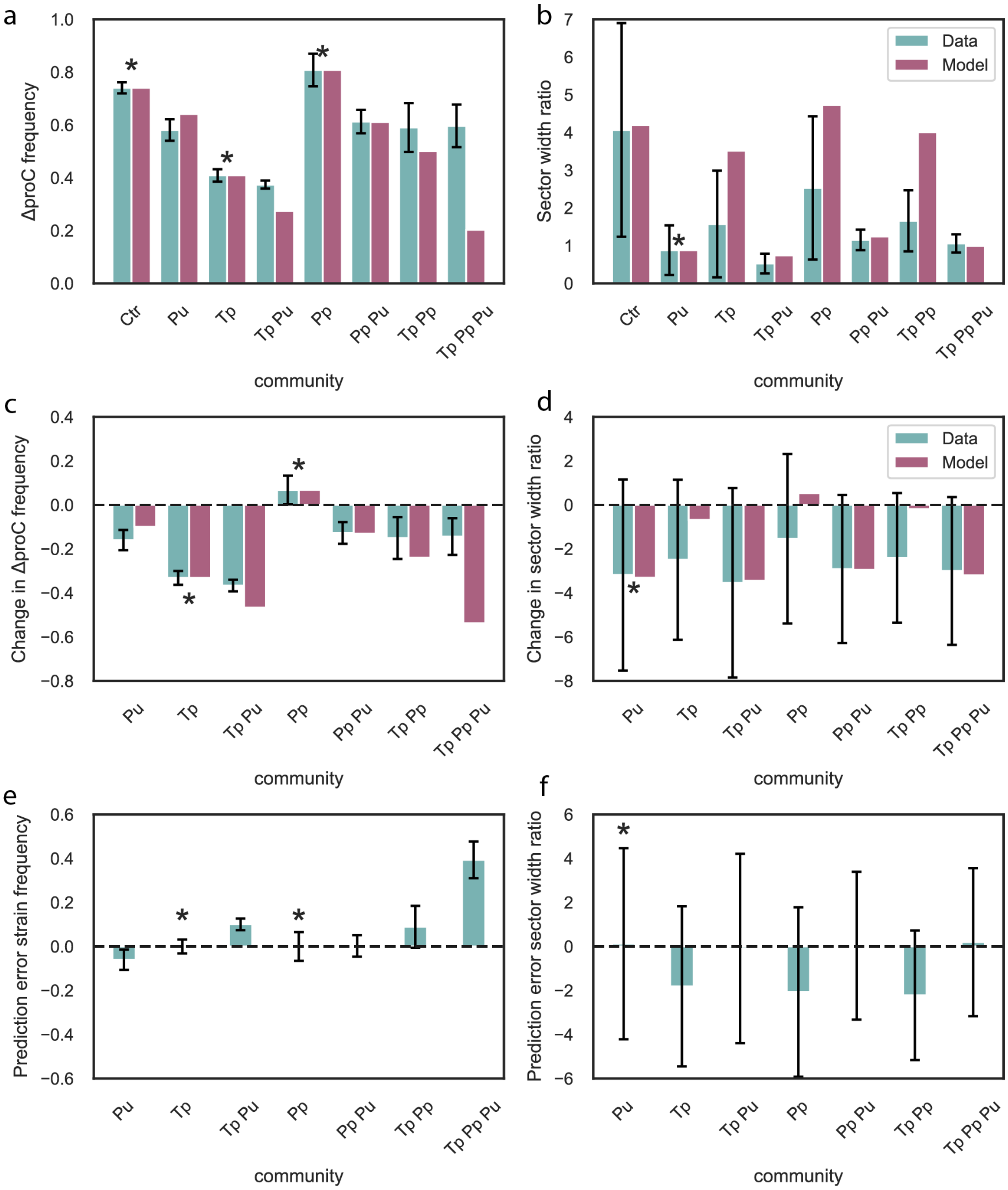


Figure S 4: **Comparison between model predictions and data.** **a** Comparison between the observed (blue) and predicted (pink) frequency of ΔproC. **b** Comparison between the observed (blue) ratio of sector widths and the predicted ratio of interaction ranges (pink). **c** Comparison between the observed (blue) and predicted (pink) change in frequency of ΔproC relative to that in the Ctr community. **d** Comparison between the change in the observed (blue) ratio of sector widths and the predicted ratio of interaction ranges (pink) relative to that in the Ctr community. **e** Prediction error for the change in the frequency of ΔproC, i.e. the difference between the blue and pink bars in panel c. **f** Prediction error for the change in sector width ratio, i.e. the difference between the blue and pink bars in panel c. **a-e** Error bars indicate 95% confidence intervals. $*$ indicates measurements which were used for parameter fitting.

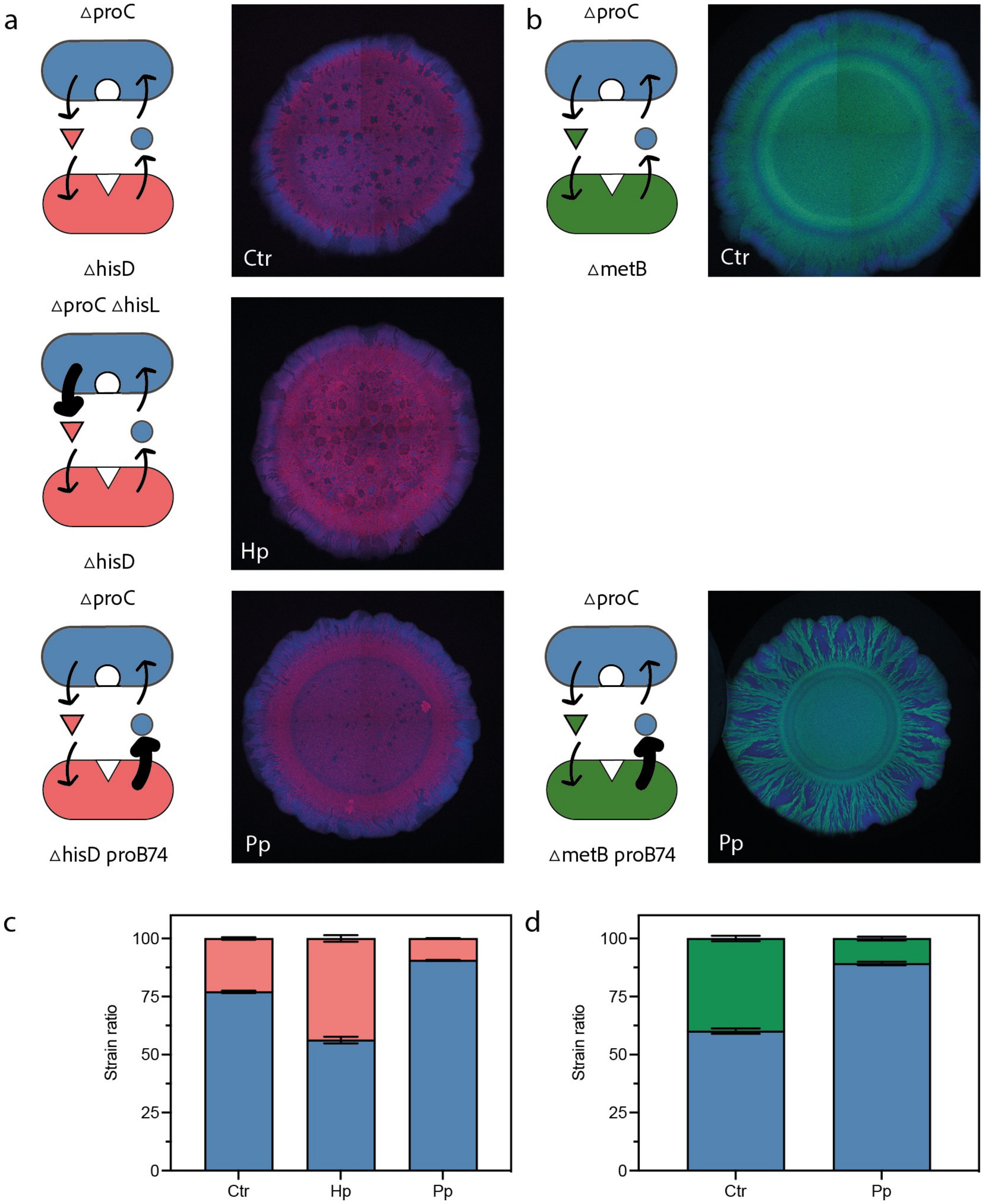


Figure S5: **Expanding to other amino acids auxotrophic strains a** Community of proline auxotroph and histidine auxotroph (Δ*hisD*). Control community (Ctr), histidine overproducer community (Hp) with Δ*hisL*, and proline overproducer community with proB74 mutant (Pp) . Representative microscopy images of the range expansion. **b** Community of proline auxotroph and methionine auxotroph (Δ*metB* ). Control community (Ctr) and proline overproducing community with *proB74* mutant (Pp). **c** Quantification of the frequencies of the two members of the proline-histidine community. Data from 3 biological replicates. **d** Quantification of the frequencies of the two members of the proline-methionine community. Data from 3 biological replicates.

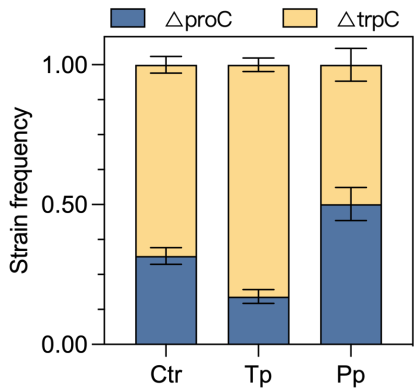


Figure S6: **Amino acid overproduction leads to a change of community composition in liquid cultures too.** Comparison between the initial community (Ctr) and communities with increased production of tryptophan (Tp) and proline (Pp) in liquid cultures (96 well plate).

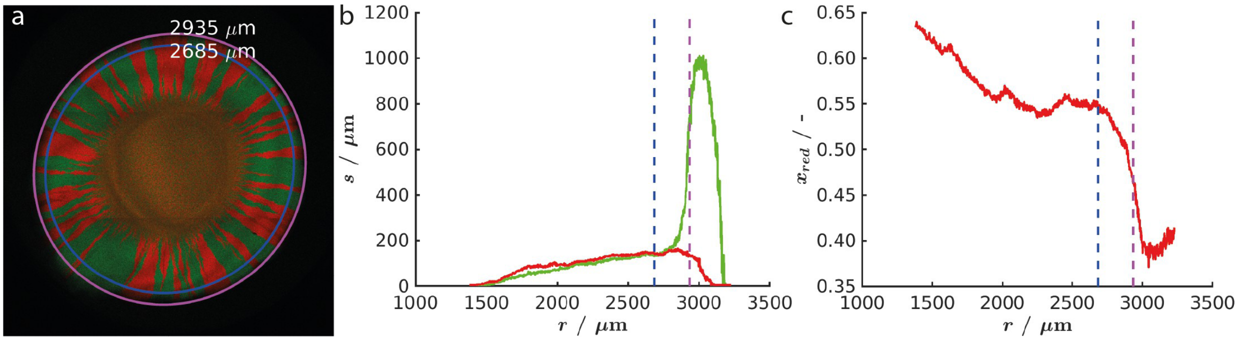


Figure S7: **The process of the image analysis. a** Image of a colony of the two labelled *E. coli* strains without auxotrophies. Magenta curve: the edge of the colony, blue curve: the radius where the separation length and the ratio of the cells were determined. The average radius of these curves were 2935 $\mu m$ and 2685 $\mu m$, respectively. **b** The separation length ($s$) as a function of the radius ($r$) for the two different cell types. **c** The ratio of the red cells ($x_{red}$) as a function of the radius. In b and c the vertical dashed coloured lines indicate the radius of the colony (magenta) and the radius where the average values were determined (blue).

**Supplementary Tables**

Table S1: Strains used in this study, derived from *E.coli* MG1655

| Strain name | Genotype | Properties | Origin |
|---|---|---|---|
| *ΔproC* | ΔproC:FRT-mCherry:FRT | Auxotrophic for proline | Dal Co et al., 2020 [1] |
| *ΔtrpC* | ΔtrpC:FRT-sfGFP | Auxotrophic for tryptophan | Dal Co et al., 2020 [1] |
| *ΔproC ΔtrpR* | ΔproC:FRT-mCherry:FRT ΔtrpR:FRT | Auxotrophic for proline and tryptophan overproducer | This study |
| *ΔtrpC proB74* | ΔtrpC:FRT - proB D107A-sfGFP:FRT | Auxotrophic for tryptophan and proline overproducer | This study |
| *ΔhisD* | ΔhisD:FRT-sfGFP | Auxotrophic for histidine | This study |
| *ΔmetB* | ΔmetB:FRT - mCherry:FRT | Auxotrophic for methionine | This study |
| *ΔmetB* | ΔmetB:FRT - sfGFP:FRT | Auxotrophic for methionine | This study |
| *ΔhisD proB74* | ΔhisD:FRT - proB D107A - sfGFP:FRT | Auxotrophic for histidine and proline overproducer | This study |
| *ΔmetB proB74* | ΔmetB:FRT - proB D107A - mCherry:FRT | Auxotrophic for methionine and proline overproducer | This study |
| *ΔmetB proB74* | ΔmetB:FRT - proB D107A - sfGFP:FRT | Auxotrophic for methionine and proline overproducer | This study |
| *ΔproC ΔhisL* | ΔproC:FRT - ΔhisL:FRT - mCherry:FRT | Auxotrophic for proline and histidine overproducer | This study |
| *ΔmetB ΔhisL* | ΔmetB:FRT - ΔhisL:FRT - mCherry:FRT | Auxotrophic for methionine and histidine overproducer | This study |
| *ΔmetB ΔhisL* | ΔmetB:FRT - ΔhisL:FRT - sfGFP:FRT | Auxotrophic for methionine and histidine overproducer | This study |

Table S2: Plasmids used in this study

| Plasmid name | Content | Origin |
|---|---|---|
| pEP28 | empty plasmid - ptet and tetR - Cm resistance | This study |
| pEP17 | *putP* under ptet promoter - tetR - CmR | This study |

Table S3: Primers used in this study

| Primer name | Sequence (5'-3') | Description |
|---|---|---|
| prEP111 | actagtagcggccgctgcagtc | Linearize pDSG360 plasmid (Forward) |
| prEP112 | ctctagtagtgctcagtatct | Linearize pDSG360 plasmid (Reverse) |
| prEP113 | agatactgagcactactagagaaagaggagaaatactag atggctattagcacacc | Amplify putP from MG1655 genome (Forward) homology to pDG360 |
| prEP114 | gactgcagcggccgctactagttaagtcccttagctttcctg | Amplify putP from MG1655 genome (Reverse) homology to pDSG360 |
| prEP167 | tacaaccgggggaggcattttgcttcccccgctaacaatggc gacatatttgtaggctggagctgcttcg | Amplify FRT Kanamycin FRT from pDK3 with homology arms for deletion of trpR (Forward) |
| prEP168 | gcattcggtgcacgatgcctgatgcgccacgtcttatcagg cctacaaaagccatggtccatatgaatatcctcc | Amplify FRT Kanamycin FRT from pDK3 with homology arms for deletion of trpR (Reverse) |
| prEP177 | taattcccatgtcagccgttaagtg | Linearize pKD4 (Reverse) |
| prEP185 | ttccatattagcacgggtcagcag | Amplify proB from MG1655 genome, includes a point mutation (A319G) on proB gene |
| prEP197 | tcccgcgcaacaaaacgccatgctttgctcgcagatggttg gcaaccgatgtaggctggagctgcttcg | Amplify FRT Kanamycin FRT from pDK3 with homology arms for deletion of proBA (Forward) |
| prEP198 | gtcaatggccttgtgaatcaaatggctacttttgcatcacccg gttttatgccatggtccatatgaatatcctcc | Amplify FRT Kanamycin FRT from pDK3 with homology arms for deletion of proBA (Reverse) |
| prEP200 | gccatggtccatatgaatatcctcc | Linearize pKD4 (Forward) |
| prEP201 | ggaggatattcatatggaccatggccgacagtcctgctaaa acgtt | Amplify proB from MG1655 genome, homology to pKD4 |
| prEP202 | cacttaacggctgacatgggaattatttacgcacgaatggtgt aatcacc | Amplify proBA from MG1655 genome, homology to pKD4 |
| prEP203 | ctgctgacccgtgctaatatggaa | Amplify proBA from MG1655 genome, includes a point mutation (A319G) on proB gene |
| prEP205 | ttcacgaacgtgaatcacggtggacaaggttaaaactaacc gggcgatgctttacgcacgaatggtgtaatcacc | Amplify FRT Kanamycin FRT and proBA (mutated) to integrate in genome |
| prEP206 | gactgcagcggccgctactagttctctagtagtgctcagtatc tctatcactga | Linearize pEP17, homology to pEP17, removing putP sequence |
| prEP207 | gccagttcaattctggtcctgccgattgagaagatgatggagtgat cgcctgtaggctggagctgcttcg | Amplify FRT Km FRT with homology arms to hisD (F) |
| prEP208 | Cgtcaggttgcggacgttttcacgcgctaaatcggtaatagtcac ggtgcgccatggtccatatgaatatcctcc | Amplify FRT Km FRT with homology arms to hisD (R) |
| prEP211 | ttactctggtgcctgacatttcaccgacaaagcccagggaacttc atcactgtaggctggagctgcttcg | Amplify FRT Km FRT with homology arms to metB (F) |
| prEP212 | tttatgcagctgacgacctttcgcccctgcctgcgcaatcacactc atttgccatggtccatatgaatatcctcc | Amplify FRT Km FRT with homology arms to metB (R) |
| prEP227 | gtggtttaggttaaaagacatcagttgaataaacattcacagaga ctttttgtaggctggagctgcttcg | Amplify FRT Km FRT with homology arms to hisL (F) |
| prEP228 | atgcaccactggaagatctgaatgtcttccagcacacatcgcctg aaagagccatggtccatatgaatatcctcc | Amplify FRT Km FRT with homology arms to hisL (R) |
| prEP250 | cattaaagaggagaaattaactatgagtgacagccagacg ctg | Amplify FRT Kanamycin FRT and proBA (mutated) |

Table S4: Parameters used for model. [a]: measured in units of the Monod constant. [b]: maximum growth rate in presence of amino acids, measured in batch cultures.

| Parameter | Description | Strains | Value | Source |
|---|---|---|---|---|
| $u_P$ | Uptake rate proline, WT | All but *putP* | 2.04 $1/s$ | [3] |
| $u_{P,putP}$ | Uptake rate proline, *putP* | *putP* | 156.2 $1/s$ | Fitted |
| $u_T$ | Uptake rate tryptophan, WT | All | 24.05 $1/s$ | [4] |
| $l_P$ | Leakage rate proline, WT | All but *proB74* | $5.29 \cdot 10^{-7}$ $1/s$ | Fitted |
| $l_{P,proB74}$ | Leakage rate proline, *proB74* | *proB74* | $2.22 \cdot 10^{-6}$ $1/s$ | Fitted |
| $l_T$ | Leakage rate tryptophan, WT | All but Δ*trpR* | $4.94 \cdot 10^{-8}$ $1/s$ | Fitted |
| $l_{T,trpR}$ | Leakage rate tryptophan, Δ*trpR* | Δ*trpR* | $1.91 \cdot 10^{-6}$ $1/s$ | Fitted |
| $D_P$ | Diffusion constant proline | All | 879 $\mu m^2/s$ | [5] |
| $D_T$ | Diffusion constant tryptophan | All | 659 $\mu m^2/s$ | [6] |
| $I_C$ | Internal concentration produced amino acid[a] | All | 20 | [1] |
| $\rho$ | Cell density | All | 0.65 | [1] |
| $L$ | Cell length | All | 5.2 $\mu m$ | [2] |
| $W$ | Cell diameter | All | 0.68 $\mu m$ | [2] |
| $\mu n$ | Maximum growth rate[b] | ΔproC | 0.24 $1/h$ | Measured |
| $\mu n$ | Maximum growth rate[b] | ΔproC *putP* | 0.049 $1/h$ | Measured |
| $\mu n$ | Maximum growth rate[b] | ΔproC Δ*trpR* | 0.29 $1/h$ | Measured |
| $\mu n$ | Maximum growth rate[b] | ΔproC *putP* Δ*trpR* | 0.048 $1/h$ | Measured |
| $\mu n$ | Maximum growth rate[b] | ΔtrpC | 0.24 $1/h$ | Measured |
| $\mu n$ | Maximum growth rate[b] | ΔtrpC *proB74* | 0.31 $1/h$ | Measured |
| $\mu n$ | Maximum growth rate[b] | WT | 0.27 $1/h$ | Measured |
